# A data acquisition and analysis pipeline for scanning angle interference microscopy

**DOI:** 10.1101/050468

**Authors:** Catherine B. Carbone, Ronald D. Vale, Nico Stuurman

**Affiliations:** Department of Cellular and Molecular Pharmacology and the Howard Hughes Medical Institute University of California San Francisco

**Author notes:** Address correspondence to Ronald Vale.

**Keywords:** Interference Microscopy, Software, Super-resolution microscopy

## Abstract

We describe open source software and hardware tools for calibrating, acquiring, and analyzing images for <u>*s*</u>canning <u>*a*</u>ngle <u>*i*</u>nterference <u>*m*</u>icroscopy (SAIM) analysis. These tools make it possible for any user with a TIRF microscope equipped with a motorized illumination unit to generate reconstructed images with nanometer precision in the axial (z) direction and diffraction-limited resolution in the lateral (xy) plane.

## MAIN

Super resolution techniques such as STORM and PALM achieve ~20 nm resolution in the lateral (xy) plane. However, achieving comparable resolution in the axial (z) direction is more difficult. Single molecule techniques (either through point spread function engineering or multi-focal imaging) usually reach ~50 nm resolution [1]. Using interference techniques (e.g. iPALM), sub-20-nm axial resolution can be obtained [2], but the complexity of the optics limits its wide use. Alternatively, surface-generated fluorescence interference contrast methods, including <u>*fl*</u>uorescence <u>*i*</u>nterference <u>*c*</u>ontrast (FLIC) microscopy or <u>*s*</u>canning <u>*a*</u>ngle <u>*i*</u>nterference <u>*m*</u>icroscopy (SAIM), are capable of nanometer resolution axial measurements [3–9]. Unlike STORM and PALM, these methods usually acquire images from large numbers of fluorophores, rather than single molecules. While utilizing relatively simple optics available on commercial microscopes, difficulties with calibration and the lack of available analysis software have hampered the widespread adoption of SAIM. Here, we describe a tool for automated angle calibration and freely available, open source software, which together create a straightforward pipeline for SAIM analysis.

In these interference contrast methods, the sample is placed on top of a transparent spacer on top of a mirror (silicon wafer with silicon oxide spacer) (**Fig. 1a**). Incident excitation light interferes with its own reflection (**Fig. 1b**), resulting in an intensity field that varies with axial distance in a manner that is theoretically predictable (**Fig. 1c, Supplementary Note**). By varying the angle of excitation, different intensity profiles can be obtained. After acquiring images at many different angles and fitting the intensities pixel by pixel to the theoretical prediction, one can determine the average height of each spatial element at nm resolution.

**Figure 1.**
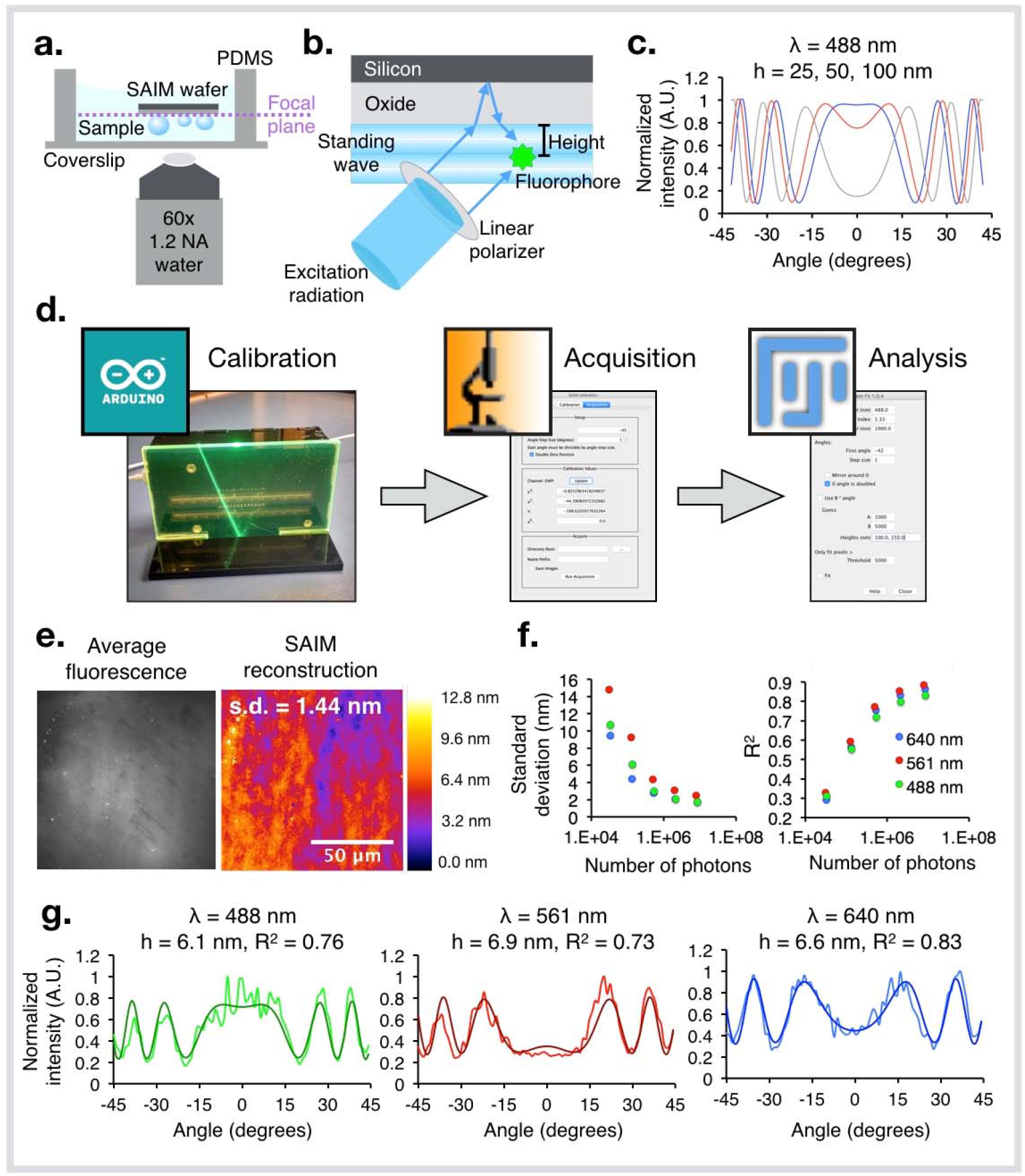
Implementation of SAIM to generate topographical maps of supported lipid bilayers. (**a**) Schematic representation of sample setup for SAIM. A sample is assembled on a silicon oxide wafer, and then the wafer is inverted, submerged in buffer for imaging. (**b**) Schematic of SAIM theory, adapted from reference [9]. Patterned illumination is achieved by interference between incident and reflected excitation light. This pattern is dependent on the angle of illumination. (**c**) Theoretical SAIM intensity profiles. Calculated angle dependent intensity profiles are shown at 25 nm (blue), 50 nm (red), or 100 nm (grey) from the surface of the oxide layer for 488 nm light with an oxide thickness of 1900 nm. (**d**) Schematic of SAIM tools. A μManager controlled Arduino device is used to calibrate the angle of laser illumination, and then a μManager plugin is used to acquire SAIM image sets. Finally these image sets are analyzed by a Fiji [14–15] plugin to output a topographical map of the sample. (**e**) SAIM height of a supported lipid bilayer labeled with DiI. Left, mean fluorescence intensity across all angles sampled. Right, SAIM reconstruction showing a height standard deviation of 1.44 nm across a 10,000 μm^2^ field. (**f**) Error in SAIM height determination at three wavelengths. Left, relation between number of photons measured per pixel and standard deviation across a 10,000 μm^2^ field. Right, relation between number of photons measured per pixel and the mean R^2^ value for the SAIM fit at each pixel across the 10,000 μm^2^ field. All measurements are shown for 488, 561, and 647 nm wavelength excitation of a supported lipid bilayer triply labeled with DiO, DiI, and DiD. (**g**) Representative single pixel fits for supported lipid bilayer measurements. Raw data shown as lighter colored line, fit shown in darker colored line.

Our SAIM pipeline begins with an improved calibration procedure (**Fig. 1d**, left). We designed a simple apparatus, controlled by a μManager plugin, that consists of an interchangeable, transparent acrylic front plate through which the laser beam travels, and two linear CCDs detecting light emerging from the front plate. The color of acrylic used for the front plate must be chosen to optimally scatter or fluoresce upon illumination by the laser being calibrated. This system defines the relation between motor position and angle of illumination by transferring the measurements to the μManager plugin and fitting with a trinomial function (**Supplementary Protocol**). Compared to manual procedures using a protractor or similar method [9–12], automated angle measurement provides a faster, more thorough calibration with sub-pixel localization of the laser position, and the ability to measure angle at finer motor steps due to the dramatic decrease in time needed for calibration. Automated calibration is especially advantageous for multicolor measurements, as it reduces the time required to calibrate each laser from 30-60 min to less than 5 min. Furthermore, this automated method is highly reproducible, which is important because errors in calibration of a few degrees can translate into errors in height measurements of greater than 10 nm (**Supplementary Fig. 1**). The building plans for this device, firmware and computer interface code are freely available (**Supplementary Protocol**).

After calibration, the sample can be analyzed by varying the angle of illumination by controlling a motorized illuminator with open source μManager software (**Fig. 1d**, middle) [13]. The acquired images then are analyzed to determine z-axis information (**Fig. 1d**, right). Our SAIM analysis software is a Fiji [14–15] plugin that: 1) plots the predicted excitation intensity at a given distance from the surface of the oxide layer as a function of angle, 2) fits the theoretical model to an experimentally determined intensity profile at a given region of interest (ROI) as a function of angle by varying estimators for height, intensity and background [9], and 3) carries out fits for all pixels in an image. The code uses an unbounded Levenberg-Marquard optimizer that minimizes the square of the difference between observed and predicted values. Parameters are restricted to physically relevant values (for instance, no negative heights are allowed). Execution speed depends on image content, though we routinely analyze every pixel in a dataset of 86 images at 1024 × 1024 pixels in 2.5 minutes on a MacBook Pro with a 2.7 GHz Intel Core i5 processor. In practice, masking images to fit only relevant pixels can accelerate processing dramatically.

Samples for SAIM are prepared on commercially available silicon substrates with an oxide spacer. Any thickness of the oxide spacer can be used, but we found that spacers of ~1900 nm provided the optimal periodicity of the SAIM function in the range of angles that could be sampled using a 1.20 NA water immersion objective (**Supplementary Fig. 2a**). In order to measure the true height of a sample by SAIM, it is critical to know the precise thickness of the oxide spacer. We measured the oxide thickness of SAIM substrates using ellipsometry (**Supplementary Fig. 2b**). Oxide spacer thickness also can be estimated using SAIM by imaging a monolayer of surface-bound fluorescent dyes of defined thickness and fitting theoretical predictions for different oxide heights to the data until the known height of the sample is derived. We used a DiO, DiI, and DiD triple labeled supported lipid bilayer (SLB) for this purpose, and fit the oxide to the known height of 6.4 nm for a phospholipid bilayer (**Supplementary Fig. 2b-c**)[16]. We recommend performing this measurement on the day of an experiment to minimize variability in measurements due to changes in the microscope and/or calibrations (**Supplementary Fig. 2d**). Once calibrated, this technique can be used to measure the SLB height with a standard deviation around one nanometer (**Fig. 1e**) at approximately 10^6^ photons per pixel (**Fig. 1f-g**).

**Figure 2.**
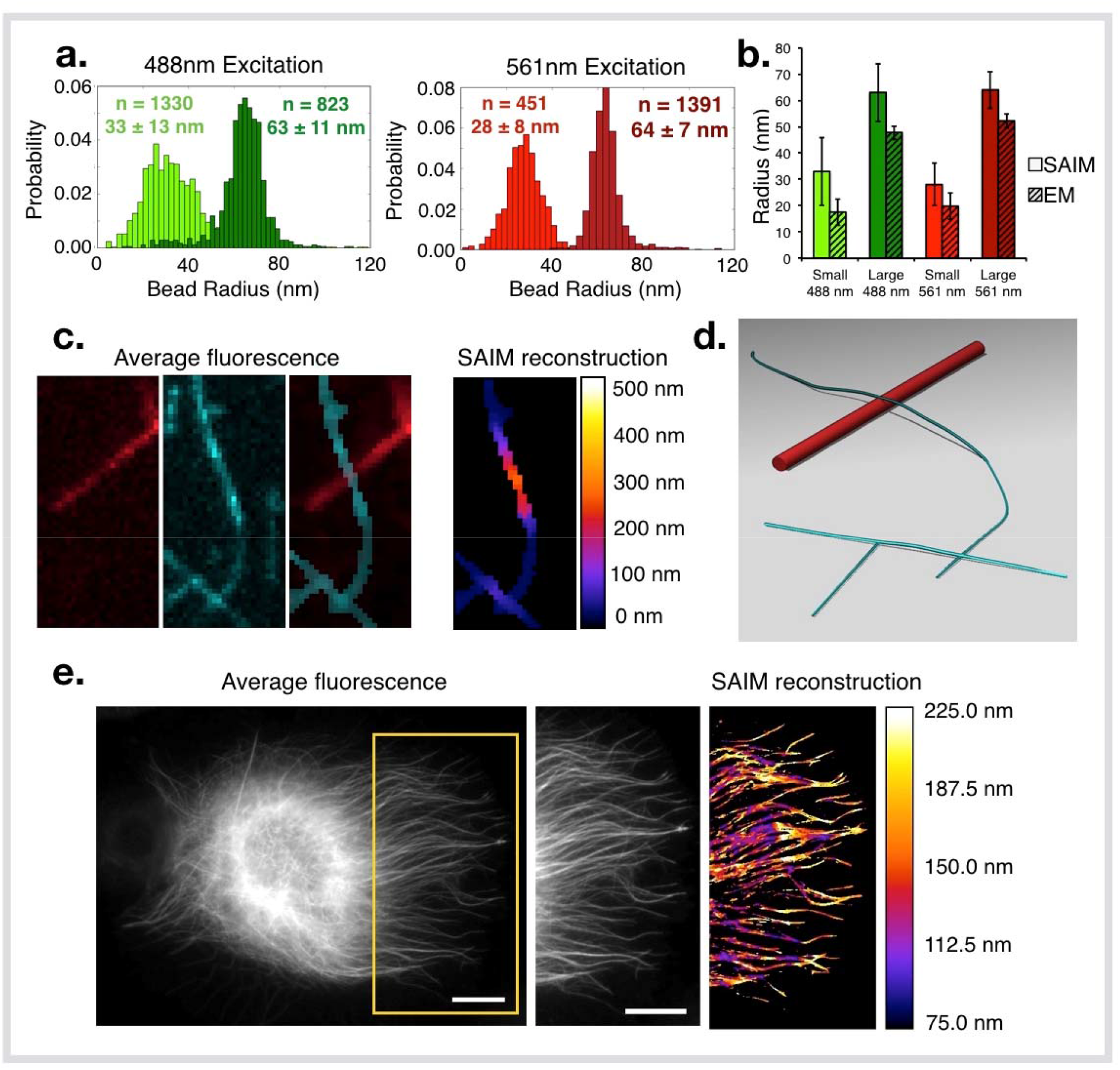
Validation of SAIM tools with experimental measurements. (**a**) Histogram of fluorescent nano-sized bead radii measured by SAIM. Left, histogram of yellow-green nanobeads. Measured heights of nominal “20 nm” radius (company provided specifications) nanobeads shown in light green (n = 1330, mean = 33 ± 13 nm), and nominal “50 nm” radius nanobeads shown in dark green (n = 823, mean = 63 ± 11 nm). Right, histogram of red nanobeads. Measured heights of nominal 20 nm radius nanobeads shown in light red (n = 451, mean = 28 ± 8 nm), and nominal 50 nm radius nanobeads shown in dark red (n = 1391, mean = 64 ± 7 nm). (**b**) Comparison of nanobead radius measured by SAIM or electron microscopy. Color-coding corresponds to panel **a**, radii measured by SAIM are solid colored, radii measured by electron microscopy are indicated by hatch marks. Error bars denote standard deviation. (**c**) SAIM height of a microtubule crossing an axoneme. Left, mean fluorescence intensity for a Cy5 labeled axoneme (red) and Alexa 647 labeled microtubule (cyan). Right, SAIM reconstruction of microtubule height showing a representative deflection at the intersection with the axoneme and intersection with other microtubules. (**d**) 3D model generated in Chimera [20] of data shown in **c**. Microtubules are shown as cylinders with a radius of 12.5 nm along spline fits through SAIM data (cyan). The axoneme is modeled as a straight cylinder with a radius of 100 nm (red). (**e**) SAIM height of microtubules in a *Drosophila* S2 cell. Left, mean fluorescence intensity for GFP-tubulin expressing S2 cell. Center, enlarged view of insert indicated by yellow box. Right, SAIM reconstruction of microtubule heights within insert.

To further test accuracy and limitations of SAIM and our analysis pipeline, we measured fluorescent nano-sized beads using both SAIM and negative stain electron microscopy. For SAIM, beads were adsorbed onto silicon substrates with an oxide spacer of ~1900 nm. Angle dependent changes in bead fluorescence intensity were fit to the optical model to obtain the axial position of the bead center. These measurements were repeated for beads with nominal 20- or 50-nm radii, and fluorescence excitation wavelengths of 488 or 561 nm (**Fig. 2a**). SAIM heights were similar to those derived from the EM measurements (**Fig. 2b**).

Next, we sought to measure nanometer scale changes in height in a biological sample. We measured the deformation of a microtubule (25 nm diameter) as it crossed on top of an axoneme (a relatively rigid, structured bundle of microtubules in a specific arrangement with a diameter of about 200 nm). The average microtubule height change at an axoneme crossing was measured by our SAIM technique to be 227 ± 44 nm (**Fig. 2c**, modeled in **Fig. 2d**). The standard deviation in the height measurements of the microtubule in areas away from crossings was 6 nm. Thus, the additional variability at axoneme crossings likely reflects heterogeneity in microtubule bending. Our automated SAIM method also is capable of measuring microtubule deformations in *Drosophila* S2 cells, as described in other cell types (**Fig. 2e**) [9,11].

In conclusion, we have developed open source tools for automated angle calibration, image acquisition and analysis methods to implement <u>*s*</u>canning <u>*a*</u>ngle <u>*i*</u>nterference <u>*m*</u>icroscopy (SAIM). With these tools, any lab with a motorized TIRF microscope supported by μManager should be able to perform SAIM experiments to measure the axial organization of fluorescently-tagged molecules at nm resolution.

## METHODS

### The SAIM calibration device

The calibration device is based on an Arduino open-source electronics platform. The device uses two linear CCDs (AMS-TAOS USA Inc., TSL-1412S) positioned above the microscope objective lens to detect fluorescence emitted by an acrylic faceplate when illuminated by an excitation laser (**Fig. 1d**, left). Based on the known, fixed vertical displacement between these two detectors the angle of that excitation light can be determined (**Supplementary Protocol**).

### The SAIM calibration plugin

This μManager [13] plugin controls the SAIM calibration device described above to execute SAIM calibration and data acquisition. This plugin consists of two parts. 1) The “Calibration” panel communicates with the calibration device to measure the relation between the refractive index corrected angle of excitation light and position of a motorized TIRF illuminator. The plugin will store calibrations for multiple channel groups that can be accessed from the acquisition panel. 2) The “Acquisition” panel reads the calibration for a given channel group, takes a user provided range of angles and the number of steps for sampling this range, and then runs a series of acquisitions at these angles.

### The SAIM analysis plugin

This Fiji [14–15] plugin consists of three parts. “SAIM Plot” plots theoretical predictions for the intensity distribution as a function of distance from the surface of the oxide layer. “SAIM Inspect” and “SAIM Fit” are very similar; however, SAIM Inspect will act on the average values of the ROI (for instance, the pixel under the cursor) and executes only a single fit, whereas SAIM Fit will analyze all pixels of the image stack. SAIM Fit will fit each pixel in the input stack and output a stack with 4 images. The first one is the height map (in nm), the second image has the r-squared values (an indication of the goodness of fit with values between 0 and 1, the closer to 1, the better the fit), the third image shows the values for “A”, a scaling parameter that accounts for variation in intensity, and the last image shows the values for “B”, an offset parameter that accounts for background fluorescence. This plugin uses the equations from the Paszek et al. paper [9] with a few extensions (**Supplementary Note**).

### Optical setup

SAIM imaging was performed on one of two Nikon TI-E microscopes equipped with a Nikon 60x Plan Apo VC 1.20 NA water immersion objective, and four laser lines (405, 488, 561, 640 nm), either a Hamamatsu Flash 4.0 or Andor iXon EM-CCD camera, and μManager software [13]. A polarizing filter was placed in the excitation laser path to polarize the light perpendicular to the plane of incidence. Angle of illumination was controlled with either a standard Nikon TIRF motorized positioner or a mirror moved by a motorized actuator (Newport, CMA-25CCCL).

### Preparation of reflective substrates with adsorbed nanobeads

N-type silicon wafers with 1900 nm ± 5% thermal oxide were purchased from Addison Engineering. Wafers were cut to approximately 1 cm^2^ using a diamond tipped pen and cleaned using air plasma for five min at a radio frequency of 18W (Harrick Plasma). 40- or 100-nm carboxylate-modified yellow-green, orange, or red fluorescent spheres (Invitrogen) were diluted in 70% ethanol, added to the wafers, and dried in a vacuum desiccator. The wafers were then washed vigorously with water, air-dried, and stored at room temperature.

### Electron microscopy

40- or 100-nm carboxylate-modified yellow-green, orange, or red fluorescent spheres (Invitrogen) were prepared by 100- to 500-fold dilution into 70% ethanol followed by sonication. To prepare grids for negative stain EM, samples were applied to freshly glow discharged carbon coated 400 mesh copper grids and blotted off. Immediately after blotting, a 2% uranyl formate solution was applied for staining and blotted off. The stain was applied five times per sample. Samples were allowed to air dry before imaging. Data were collected on a Tecnai T12 microscope (FEI) equipped with a 4K × 4K CCD camera (UltraScan 4000, Gatan). 100 nm nanobeads were imaged with a pixel size of 0.58 pixels per 1 nm and magnification of 6,500x, and 40 nm nanobeads were imaged with a pixel size of 0.98 pixels per 1 nm magnification of 11,000x.

### Preparation of reflective substrates with supported lipid bilayers

Silicon wafers with 1900 nm oxide spacers were obtained from Addison Engineering, cut, and cleaned using the same process as for for nanobead imaging. Synthetic 1,2-dioleoyl-*sn*-glycero-3-phosphocholine (POPC; Avanti, 850457), 1,2-dioleoyl-*sn*-glycero-3-[(N-(5-amino-1-carboxypentyl)iminodiacetic acid)succinyl] (nickel salt, DGS-NTA-Ni; Avanti, 790404) and 1,2-dioleoyl-*sn*-glycero-3-phosphoethanolamine-N-[methoxy(polyethylene glycol)-5000] (ammonium salt, PEG5000-PE; Avanti, 880220) were purchased from Avanti Polar Lipids. Small unilamellar vesicles (SUVs) were prepared from a mixture of 95.5% POPC, 2% DGS-NGA-Ni, and 0.5% PEG5000-PE. The lipid mixture in chloroform was evaporated under argon and further dried under vacuum. The mixture was then rehydrated with phosphate buffered saline pH 7.4 and cycled between −80°C and 37°C 20 times, and then centrifuged for 45 min at 35,000 RCF. SUVs made by this method were stored at 4°C and used within two weeks of formation. To make labeled supported lipid bilayers, wafers were submerged in PBS in freshly plasma cleaned custom PDMS chambers on RCA cleaned glass coverslips. 100 µl of SUV solution containing 0.5 to 1 mg/ml lipid was added to the coverslips and incubated for 30 min. Unadsorbed vesicles were removed by washing three times with PBS, then bilayers were stained for 20 min with approximately 100 ng/mL DiO, DiI and/or DiD solution in PBS (Invitrogen). Wafers were again washed three times with PBS prior to imaging.

### Preparation of microtubules

Tubulin was purified from pig brain, and biotinylated or fluorescently labeled tubulin were prepared as described [17]. A mixture of unlabeled tubulin, biotin-tubulin, and fluorescent tubulin (~10:1:1 ratio) was assembled in BRB80 (80 mM PIPES, 1 mM EGTA, 1 mM MgCl_2_) + 1 mM GTP for 15 min at 37°C and polymerized MTs were stabilized with 20 µM taxol (Sigma, T1912). MTs were centrifuged over a 25% sucrose cushion to remove aggregates and unassembled tubulin before use.

### Assembly of microtubule-axoneme crossings on reflective substrates

Purified, fluorescently labeled sea urchin axonemes [18] were flowed onto silicon wafers and allowed to adhere for 10 min. After washing excess unbound axonemes using BRB80 buffer (80 mM Pipes pH 6.8, 2 mM MgCl_2_, 1 mM EGTA), the chip was coated twice with 5 mg/ml BSA-biotin (Thermo Scientific, 29130), washed with BRB80, coated with 0.5 mg/ml streptavidin (Vector Labs, SA-5000), and washed again with BRB80 plus 10 µM taxol. Polymerized microtubules were then added to the wafer and allowed to adhere for 5-10 min. Unbound microtubules were washed away using BRB80/taxol, and the wafer was then submerged in BRB80/10 µM taxol with an oxygen scavenging system [19] for imaging.

### Cell culture and sample preparation

Drosophila S2 cells expressing GFP-tubulin were cultured in Schneider’s Drosophila Medium with 10% fetal bovine serum and Antibiotic-Antimycotic (ThermoFisher Scientific). Silicon wafers with 1900 nm oxide spacers were cleaned using the same process as for nanobead imaging. Wafers were then incubated with 0.5 mg/mL Concanavalin A (Sigma Aldrich, C-7275) and dried under ultraviolet light. Cells were added to wafers and allowed to settle overnight. To fix, wafers were incubated in 0.25% glutaraldehye, 4% paraformaldehyde and 0.5% Triton X-100 in phosphate buffered saline for 10 min. Glutaraldehyde induced auto-fluorescence was quenched by a 7 min incubation in 1 mg/ml NaBH4 in phosphate buffered saline. Samples were stored at 4°C in 2% BSA, 0.1% Triton X-100, 0.1% NaN_3_ in phosphate buffered saline.

## SOFTWARE AVAILABILTY

SAIM analysis and μManager acquisition software are available under the Berkeley Software Distribution (BSD) license. Full documentation and examples are available at the project pages http://imagej.net/Saim and https://valelab.ucsf.edu/~kcarbone/SAIM. Development is hosted on GitHub at https://github.com/nicost/saimAnalysis/ and https://github.com/kcarbone/SAIM_calibration.

## AUTHOR CONTRIBUTIONS

C.B.C., R.D.V., and N.S. designed the research. N.S. and C.B.C. developed the software and calibration device; C.B.C. collected data to test software, C.B.C. and N.S. drafted the article; C.B.C., R.D.V., and N.S. edited the article; and all authors read and approved the final article.

## ACKNOWLEDGMENTS

We would like to thank Gira Bhabha and Stefan Niekamp for help with electron microscopy, axoneme and microtubule experiments and Jongmin Sung and Sonny Vo (director of the Leia3d Advance Lithography Center in Palo Alto, CA) for ellipsometry measurements. We thank Tom Goddard for help generating a model of the microtubule/axoneme crossing data. We are grateful to the Valerie Weaver and Matt Rubashkin for initial help implementing SAIM and to Matt Paszek and Bill Shin for useful discussions about the SAIM software and calibration device. We thank Gira Bhabha and Ankur Jain for comments on the manuscript. The authors acknowledge funding from the National Institutes of Health (R01EB007187) and the Howard Hughes Medical Institute.

